# Gut Analysis Toolbox: Automating quantitative analysis of enteric neurons

**DOI:** 10.1101/2024.01.17.576140

**Authors:** Luke Sorensen, Adam Humenick, Sabrina S.B. Poon, Myat Noe Han, Narges Sadat Mahdavian, Ryan Hamnett, Estibaliz Gómez-de-Mariscal, Peter H. Neckel, Ayame Saito, Keith Mutunduwe, Christie Glennan, Robert Haase, Rachel M. McQuade, Jaime P.P. Foong, Simon J.H. Brookes, Julia A. Kaltschmidt, Arrate Muñoz-Barrutia, Sebastian K. King, Nicholas A. Veldhuis, Simona E. Carbone, Daniel P. Poole, Pradeep Rajasekhar

## Abstract

The enteric nervous system (ENS) plays an important role in coordinating gut function. The ENS consists of an extensive network of neurons and glial cells within the wall of the gastrointestinal tract. Alterations in neuronal distribution, function, and type are strongly associated with enteric neuropathies and gastrointestinal (GI) dysfunction and can serve as biomarkers for disease. However, current methods for assessing neuronal counts and distribution suffer from undersampling. This is partly due to challenges associated with imaging and analyzing large tissue areas, and operator bias due to manual analysis. Here, we present the Gut Analysis Toolbox (GAT), an image analysis tool designed for characterization of enteric neurons and their neurochemical coding using 2D images of GI wholemount preparations. GAT is developed for the Fiji distribution of ImageJ. It has a user-friendly interface and offers rapid and accurate cell segmentation. Custom deep learning (DL) based cell segmentation models were developed using StarDist. GAT also includes a ganglion segmentation model which was developed using deepImageJ. In addition, GAT allows importing of segmentation generated by other software. DL models have been trained using ZeroCostDL4Mic on diverse datasets sourced from different laboratories. This captures the variability associated with differences in animal species, image acquisition parameters, and sample preparation across research groups. We demonstrate the robustness of the cell segmentation DL models by comparing them against the state-of-the-art cell segmentation software, Cellpose. To quantify neuronal distribution GAT applies proximal neighbor-based spatial analysis. We demonstrate how the proximal neighbor analysis can reveal differences in cellular distribution across gut regions using a published dataset. In summary, GAT provides an easy-to-use toolbox to streamline routine image analysis tasks in ENS research. GAT enhances throughput allowing unbiased analysis of larger tissue areas, multiple neuronal markers and numerous samples rapidly.

## Introduction

The enteric nervous system (ENS) is a network of neurons and glial cells located within the wall of the gastrointestinal (GI) tract. The ENS extends along the esophagus to the rectum and is estimated to be comprised of ∼168 million neurons, which is comparable to the numbers in the spinal cord in humans, mice, and guinea pigs (Michel et al. 2022a). It is critical for the regulation of secretion, absorption, and immune function, and for coordination of gut motility (Furness 2012). The absence or loss of enteric neurons results in GI dysfunction as evidenced in enteric neuropathies such as Hirschsprung disease, achalasia, and Chagas disease (Burns et al. 2016; Heuckeroth 2018; Schäppi et al. 2013; Vaezi et al. 2016). Alterations to specific enteric neuron populations that express distinct combinations of neuropeptides, enzymes or neurochemicals are also evident in other diseases that impact gut function. These include inflammatory bowel disease (Brierley & Linden 2014), diabetes (Chandrasekharan et al. 2011; Ingrid et al. 2013), and Alzheimer’s disease (Niesler et al. 2021; Semar et al. 2013; Van Ginneken et al. 2011). Researchers use enteric neuronal counts as a key metric to describe any neurochemical changes in these diseases. Typically, this is achieved by manually counting cells in small intestinal segments, or more recently, via semi-automated methods (Cairns et al. 2021; Kapur 2013; Kobayashi et al. 2021; Nestor-Kalinoski et al. 2022; Schäppi et al. 2013). These cell counts from localized areas within a specimen are then used to make inferences about broader changes to the bowel region being studied and any changes associated with disease. However, the estimated number of cells counted can be affected by:

- Tissue preparation examined: Cell density estimates using tissue sections can be prone to sampling and operator bias compared to the use of wholemounts (Kapur 2013; Swaminathan & Kapur 2010).
- Age of the animal (Gamage et al. 2013).
- Tissue region examined: Regional differences in the distribution of ENS circuitry (Hamnett, Dershowitz, Sampathkumar, et al. 2022; Nestor-Kalinoski et al. 2022).
- Sampling: The number of tissue specimens and locations sampled (Nestor-Kalinoski et al. 2022).
- Operator bias: Bias during the tissue preparation, sampling, or manual counting steps (Kapur 2013; Schäppi et al. 2013).

The major limitation of current approaches for neuronal quantification in large tissue specimens is the use of manual cell counting. This process is slow, labor intensive, and prone to inter-observer variability. In our opinion, the continued use of manual counting processes is largely due to the lack of easy-to-use, ENS-specific image analysis software.

The need for image analysis software in neurogastroenterology is evidenced by the increasing number of image analysis workflows and tools becoming available in recent years, such as COUNTEN and the use of machine learning approaches in Fiji (Cairns et al. 2021; Kobayashi et al. 2021). However, to use these workflows computational expertise is required, and the software parameters need to be optimized for new data sets. COUNTEN is highly dependent on images of robust and homogeneous staining, which is not always achievable for all intestinal preparations (Kobayashi et al. 2021). Multiple variables can influence image quality, such as low signal-to-noise ratio, variations in sample preparation, quality of dissection of layers which can affect antibody penetration and visualization of the ENS, and the specific antibodies, markers, and fluorophores used during staining. Furthermore, image acquisition specifications, including the bit-depth and dynamic range, can significantly affect downstream image analysis. All these factors can pose challenges to the widespread adoption of such software, necessitating the development of customized analytical workflows for each use case.

We developed the Gut Analysis Toolbox (GAT) for the Fiji distribution of ImageJ (Schindelin et al. 2012). GAT can be used to analyze and quantify cells within the ENS. GAT uses deep learning-based (DL) cell segmentation models developed with StarDist (Schmidt et al. 2018) for segmenting enteric neurons and neuronal subtypes. A pre-trained TensorFlow model was used to segment ganglia, which is accessible in Fiji using deepImageJ (Gómez-de-Mariscal et al. 2021). These models are integrated into GAT for rapid and reproducible quantification of key metrics such as total neuronal counts, neurochemical marker distribution, and cell number per ganglion (**Fig. 1**). The DL models were trained on manually annotated data from mouse, rat, and human colon wholemounts to ensure they effectively segmented a wide variety of images (Chen et al. 2023; DiCello et al. 2020; Graham et al. 2020b; Nestor-Kalinoski et al. 2022). Images were acquired using confocal and widefield microscopes from different research groups. Proximal neighbor analysis was used to characterize neuronal distribution using CLIJ (Haase, Jain, et al. 2020; Haase, Royer, et al. 2020). Comprehensive installation and usage instructions, along with sample data and tutorial videos, can be found in the documentation available at https://gut-analysis-toolbox.gitbook.io/docs/. The GAT workflow is also usable from within the QuPath software to enable analysis of large 2D images (Bankhead et al. 2017). The enhanced throughput of GAT facilitates sampling and analysis of larger tissue areas, thus minimizing potential sampling errors and biases.

**Figure 1:**
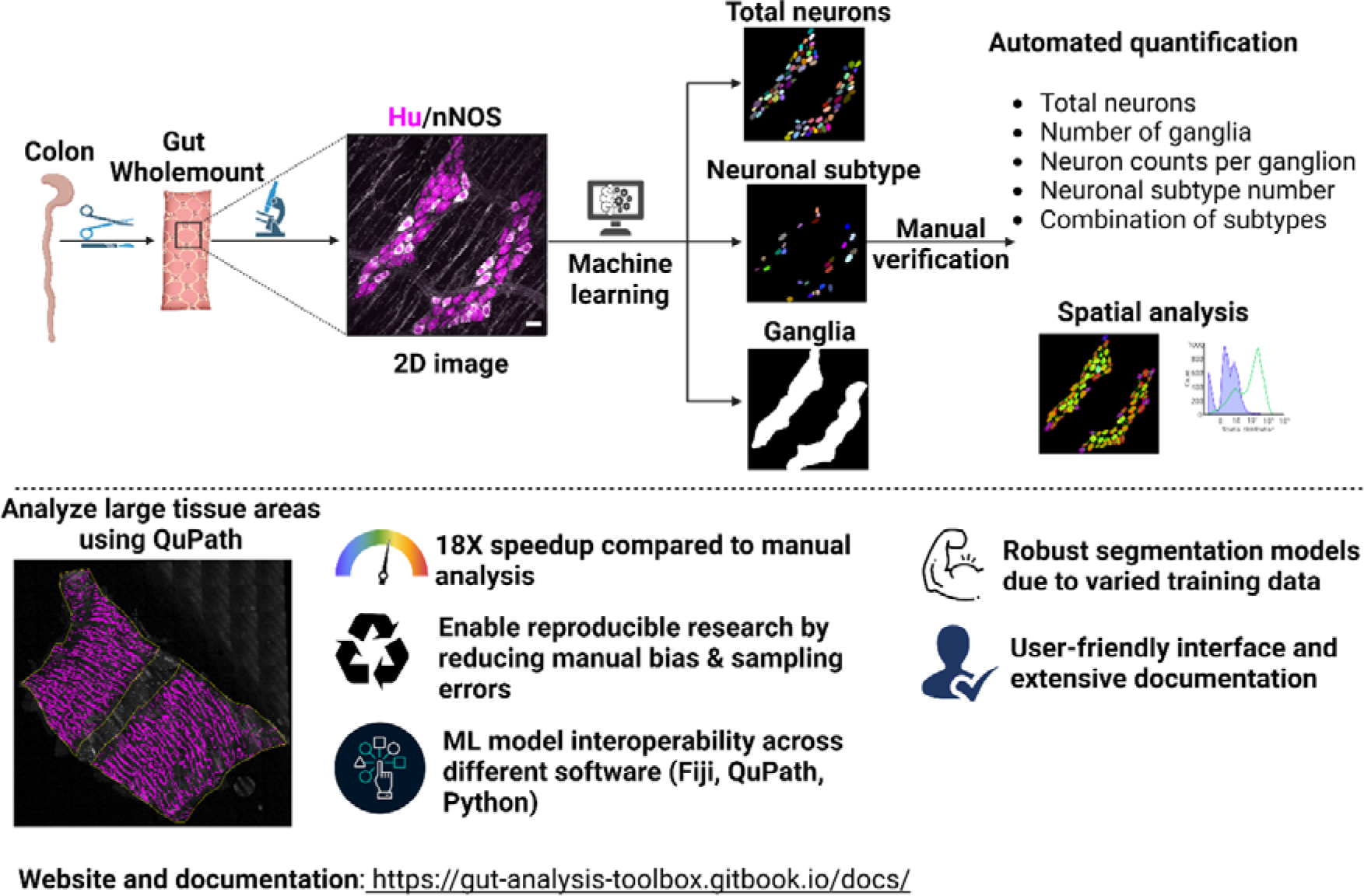
Overview of GAT workflow. GAT can segment neurons, neurons expressing neurochemical markers, and ganglia in fluorescently labeled 2D images using pre-trained DL models. GAT allows manual verification of the segmentation, followed by automated quantification of cell counts. The cellular distribution can be subsequently quantified via proximal neighbor analysis. Large 2D images can be analyzed in QuPath using the GAT DL models. Extensive documentation and videos on how to use GAT are available at: https://gut-analysis-toolbox.gitbook.io/docs/. Myenteric wholemount images from DiCello et al. (2020) and Howard (2021) (Figure created using Biorender.com).

## Results

To develop effective deep learning models, it is essential to have diverse datasets that encompass the inherent variability in images from various sources. To meet this need, we collected ENS images from four different research laboratories (Chen et al. 2023; DiCello et al. 2020; McQuade et al. 2021; Poon et al. 2022) and two publicly available datasets (Graham et al. 2020a; Howard 2021). Images were captured using a variety of microscopes, and the labeled tissues originated from different animal species, including mice, humans, and rats, as detailed in **Table S1.** The details for images with pan-neuronal marker Hu are summarized in **Table S1**. For the enteric neuron subtype model (**Table S2**), nine neurochemical markers were used, with 42% of the images representing nNOS. The ganglia model was trained using various neuronal markers in combination with Hu, as listed in **Table S3**. The effects of sample preparation and inadequate sampling on enteric neuron counts were also tested, see the supplementary information section (**Figures S1, S2** and **Supp. Text**).

### Segmentation models

StarDist (Schmidt et al. 2018) was used to train DL models for segmenting all enteric neurons. A U-Net architecture was utilized for the ganglia model (Ronneberger, Fischer & Brox 2015). The training and evaluation of the models were conducted using ZeroCostDL4Mic (von Chamier et al. 2021). Further details on the curation of training data and software versions used can be found in the methods and supplementary methods section. The ZeroCostDL4Mic Google Colab notebooks used for training and quality control are attached as supplementary files, and available online (https://zenodo.org/records/6096664).

The performance of the ‘enteric neuron model’ was evaluated on a test dataset, and compared to a widely used cell segmentation software, Cellpose (v0.7) (Stringer et al. 2021). **Fig. 2A** presents the evaluation of object detection accuracy (F1 score) and shape alignment accuracy (intersection over union, IoU) (Caicedo et al. 2019). A higher F1 score at a higher IoU indicates better segmentation performance (**Fig. 2A**). The Hu segmentation results showed comparable performance between the StarDist and Cellpose (cyto2) models. However, when examining the neuronal subtype models, a rightward shift in F1 scores of the StarDist model indicates a modest improvement when compared to the Cellpose cyto2 model (**Fig. 2A**). As StarDist approximates the shape of a cell using star-convex polygons, the predicted objects have smoother outlines relative to the original cell shape (**Fig. S5).**

**Figure 2.**
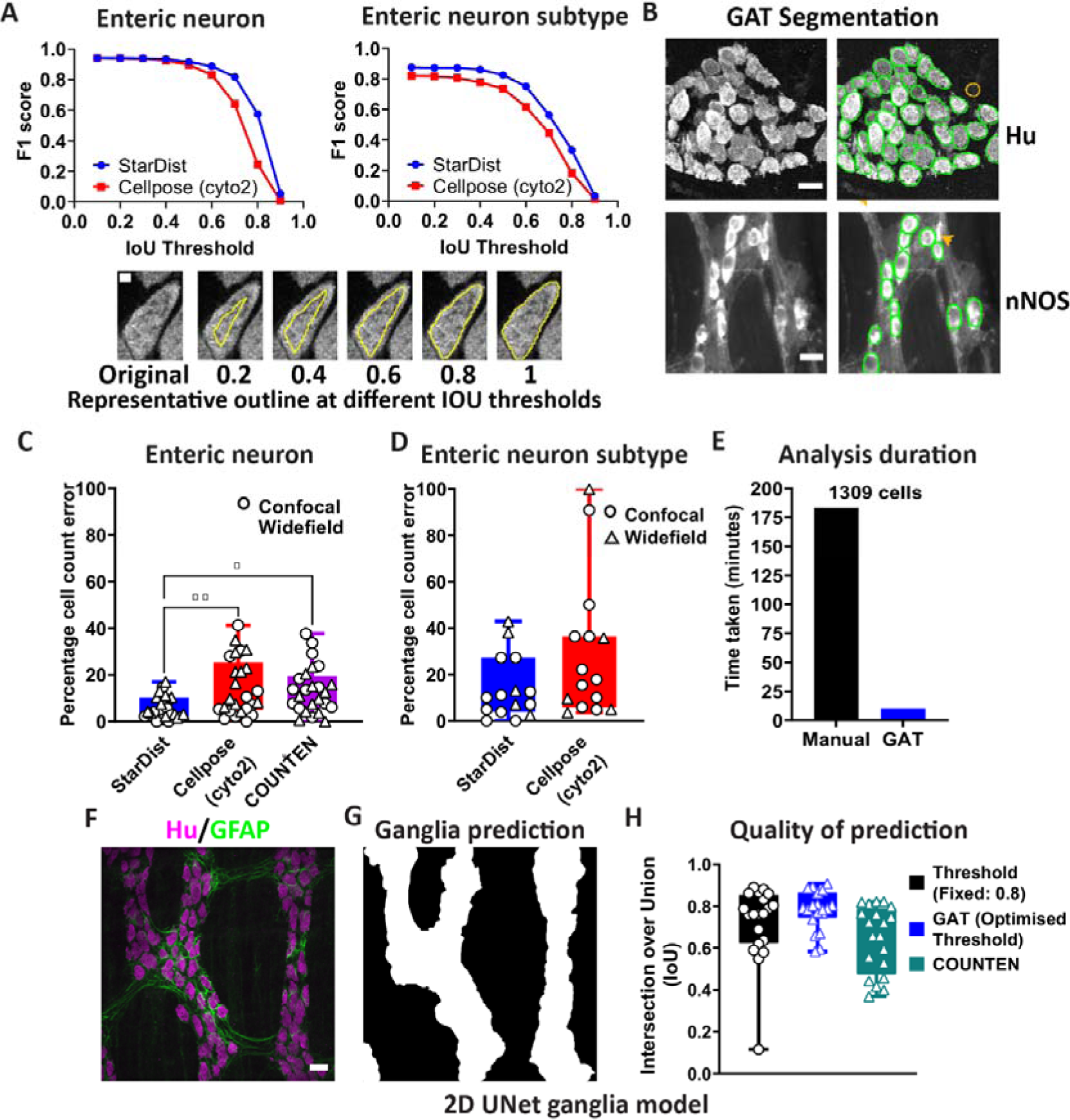
GAT segments neurons and ganglia with high accuracy. **(A)** Segmentation of Hu labeled enteric neurons by the GAT StarDist model (**Table S1**) and Cellpose (cyto2 model) had a comparable F1-score. However, the GAT StarDist model for other neurochemical markers (**Table S2**) had a better F1 score at various Intersection Over Union (IoU) thresholds. Representative images of different IoU thresholds are depicted as cell outlines below the graph (Scale bar = 5 µm). **(B)** The StarDist model could be used to detect bright versus dim cells and overlapping cells. Green outlines highlight GAT-detected cells. The orange outline shows a false positive cell, and the orange arrowhead indicates a missed cell (scale bar = 20 µm; pixel size of 0.5µm/pixel was used for segmentation). **(C)** When used to count enteric neurons, the lowest percentage of error was achieved using GAT StarDist model compared to Cellpose models and COUNTEN (20 images, 3830 cells). **(D)** The StarDist enteric neuron subtype model produced a lower percentage error compared to Cellpose when used to count different enteric neurons based on the neurochemical marker expressed (15 images, 359 cells). **(E)** The total time taken to segment 1309 neurons was significantly faster using GAT compared to manual segmentation. **(F, G)** A 2D U-Net model was trained to segment ganglia based on the presence of various markers (**Table S3**) that label the neuron or glial fibers. The representative image demonstrates prediction and segmentation using Hu and GFAP. **(H)** The ganglia segmentation model has a mean IoU of 0.72 ± 0.17 when using a fixed threshold of 0.8, and 0.78 ± 0.09 with a deepImageJ optimized threshold for each image. COUNTEN had a lower IoU of 0.645 ± 0.16 (n = 20 images, scale bar = 30 µm).

The primary goal of GAT is to estimate cell counts and this metric is not necessarily captured by the F1 score which evaluates segmentation quality. The StarDist models approximate cell shape which leads to smoother outlines, meaning the final segmentation result will slightly differ from the ground truth leading to reduced F1 scores (**Fig. S5)**. The ‘percentage cell count error’, which is the ‘(predicted number - ground truth number) /ground truth number x 100’, is a more direct measure of the accuracy of these DL models for evaluating cell count, compared to the F1 score. Importantly, this approach allows for an objective comparison with COUNTEN using the default settings recommended by the authors (Kobayashi et al. 2021). In this context, a lower percentage error indicates enhanced performance. The percentage cell count error was estimated using the same test datasets as those used for the F1 score. The StarDist neuron model had significantly better accuracy with only 6.09 ± 4.8 % error in cell counts, compared to 14.6 ± 12.1 % for Cellpose (20 images, 3830 neurons, Mean ± SD, one-way ANOVA). The percentage error for COUNTEN (**Fig. 2C**) was also higher than GAT at 13.54 ± 10 %. When testing the accuracy of segmenting enteric neurons expressing the markers calbindin (Calb), calretinin (Calret), neuronalNOS (nNOS) and neurofilament M (NFM), the StarDist model demonstrated a similar cell count percentage error (13.8 ± 13.1%) compared to Cellpose (29.5 ± 29.3 %) (**Fig. 2D**). Notably, being a generalist cell segmentation algorithm, Cellpose still had high IoU scores on the enteric neuron segmentation task, with higher cell count accuracies for 2D confocal images (**Fig. 2C**). The Cellpose predictions had higher cell splitting and merging errors (Wolny et al. 2020) compared to StarDist (**Fig. S6**). Training the Cellpose cyto2 model on these enteric neuron datasets will most likely produce a better performing Cellpose enteric neuronal model (Pachitariu & Stringer 2022). However, this was not explored due to the lack of a standalone Fiji plugin for Cellpose.

Semi-automation using GAT increased analysis throughput. Manual segmentation of 1309 enteric neurons took 183 minutes in total, across 3 researchers, whereas segmentation using GAT and the enteric neuron DL model took only 10.4 minutes by a single person (**Fig. 2E**).

To enable segmentation of ganglia using GAT, a 2D U-Net model was trained using ZeroCostDL4Mic(von Chamier et al. 2021) on images of Hu labelling in combination with a second marker for neuronal or glial fibers (**Table S3)**. The ganglia model was subsequently exported in a format compatible for use within the Fiji deepImageJ plugin (Gómez-de-Mariscal et al. 2021). A limitation is that the postprocessing threshold value applied to the probability map output from deepImageJ can impact the accuracy of the ganglia outline. To evaluate this an arbitrary threshold of 0.8 was compared with the default deepImageJ optimized threshold. The GAT ganglia model performed better compared to COUNTEN when measuring the IoU, particularly when using the optimized threshold (**Fig. 2F**) (GAT: IoU of 0.78 ± 0.09 for optimized threshold; 0.72 ± 0.17 for a fixed threshold of 0.8, vs COUNTEN: 0.64 ± 0.16, Mean ± SD). Not all laboratories use markers to label the ganglionic border. In this instance, the neuronal outline can be expanded by a user-specified distance to approximate ganglionic area.

### Proximal neighbor analysis in GAT can detect regional differences in cellular distribution

An accurate estimation of neuronal densities in gut wholemounts can be impacted by the degree of stretch applied during tissue processing. Nonetheless, the stretch applied does not alter the cellular architecture and spatial relationships between cells. Any changes in cell density results in a proportional change in the number of neighbors around each cell. As the spatial relationship is unaffected by tissue stretch, the number of proximal neighbors (PNs) is a robust measure for characterizing cellular distribution. To apply PN measurements in GAT, a threshold distance value was determined to define the distance within which cells were considered neighbors. The edge-to-edge distances between the segmented neurons in the ganglia were measured in images of the myenteric wholemounts of the mouse colon (**Table S1, & S3**). To define a threshold distance value for considering a cell a PN, the average PN distance between neurons (edge to edge distance) in the ganglia was calculated using the ‘Local Thickness’ plugin (Dougherty & Kunzelmann 2007) in Fiji (**Fig. S7**). For the myenteric wholemounts of the mouse colon, this was determined to be 6.32 ± 5.17 µm (n=3643 cells, 130 ganglia, Mean ± S.D., N=11), which was rounded up to 6.5 µm for use within GAT (**Fig. S7 C** & **D).** This value is customizable in GAT to account for differences in tissue preparation or varying cell sizes across species. The neighbor count map allowed the examination of any changes associated with different distance values.

To evaluate the robustness of spatial analysis with uneven tissue stretch, an image of the same region of a myenteric wholemount of the Wnt1-cre:Rosa26-tdTomato mouse colon was acquired under poorly stretched and stretched conditions (**Fig. 3A**). Stretch led to a ∼55.3 % increase in the tissue area examined (152318 µm^2^ vs. 245652 µm^2^ for unstretched), total ganglionic area (26573 µm^2^ vs. 44844 µm^2^) and cell densities (2696 neurons/mm^2^ vs. 1493 neurons/mm^2^, **Fig. 3A**). Using the PN analysis, no significant difference was found between the average PNs around each neuron for both stretched and unstretched tissue 2.76 ± 1.25 vs 2.98± 1.21 (Mean ± S.D., p=0.28 Students t-test, N=1). Importantly, the distribution of PNs for each cell was similar for both stretched and unstretched tissue (**Fig. 3C**), showing that GAT can capture the underlying cellular distribution even with significant differences in tissue area.

**Fig. 3:**
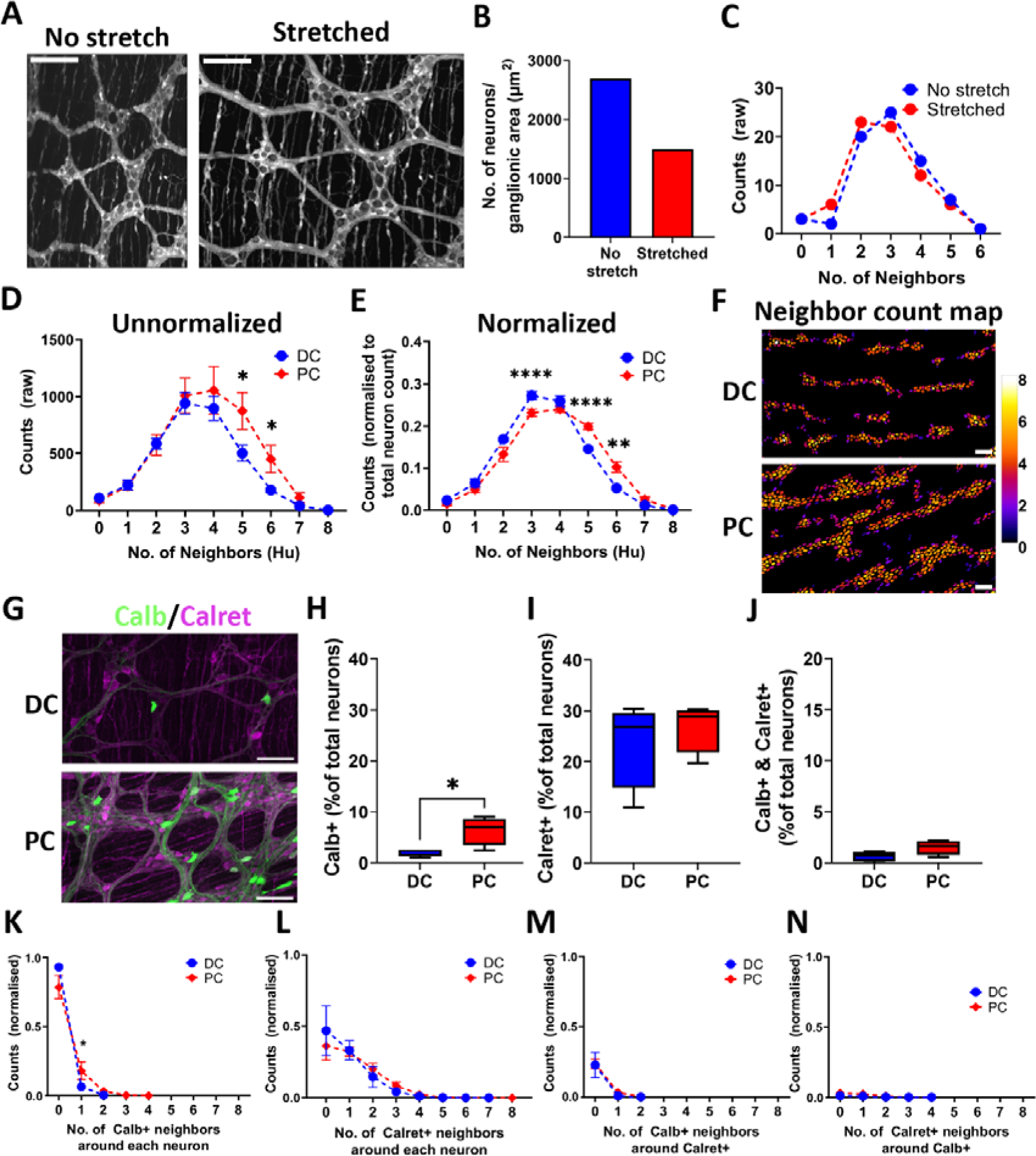
Spatial analysis in GAT can adjust for non-uniform tissue stretching and can objectively describe region-specific differences in neuronal distribution. **(A)** Both panels show the same field of view (FOV) of a myenteric plexus wholemount preparation of the Wnt1-cre:Rosa26-tdTomato mouse colon. The first panel shows the specimen that was pinned in a petri dish with ‘no stretch’ and the second panel is in the presence of ‘stretch’. Stretching the specimen leads to altered calculations of neuronal density (2696 neurons/mm^2^ vs 1493 neurons/mm^2^) **(B)**. **(C)** ‘Proximal neighbor’ (PN) analysis shows similar distribution results for non-stretched versus stretched preparations. **(D)** Proximal neighbor distribution for the proximal colon (PC) vs distal colon (DC) without normalization to the total cell count shows that PC has larger neuronal clusters with with significantly larger neuronal clusters at neighbor counts of 5 and 6. **(E)** Normalizing for total neuron count (Hu+ neurons) revealed that the DC had significantly smaller neuronal clusters at neighbor count of 3, whereas PC had significantly larger neuronal clusters at neighbor counts of 5 and 6. The difference in number of neighbors across regions which can be visualized in a neighbor count map **(F)**. **(G, H)** Calb+ positive neurons were significantly greater in proportion in the PC compared to DC, whereas there was no significant difference in the distribution of Calret+ neurons **(I)** or Calret+/ Calb+ double-positive neurons **(J)**. This was reflected in the normalized PN distribution plots, where a significantly higher proportion of neurons had one Calb+ neighbor in the PC relative to the DC. No regional differences in Calret+ neurons **(K)** or preferential association between Calret+ and Calb+ neurons were detected **(L, M, N)** (Scale bar = 100 µm).

Spatial analysis in GAT enables the quantification of cellular distribution and can be used to evaluate differences across gut regions or in different disease states. To illustrate this objectively, images of the proximal colon (PC) and distal colon (DC) were analyzed from a published dataset (Hamnett, Dershowitz, Gomez-Frittelli, et al. 2022; Hamnett, Dershowitz, Sampathkumar, et al. 2022). Others have demonstrated that the PC has larger ganglia and greater neuronal density in comparison to the DC (Li et al. 2019; Nestor-Kalinoski et al. 2022). This was confirmed by our analysis using GAT (neurons per ganglion: 20.7 ± 30.3 for DC vs 56.5 ± 202.3 for PC, Mean ± S.D. respectively, unpaired t-test with Welch’s correction, p<0.0001, n=4). The large S.D. values are reflective of the variations in ganglia sizes for each region. Neurons in the PC had greater number of neighbors than those in the DC (Fig. 3D; raw counts: 503.8 ± 68.6 vs. 874.8 ± 161 for 5 neighbors, 180.3 ± 20.4 vs. 451.5 ± 120.2 for 6 neighbors for DC vs. PC respectively, Mean ± S.D., unpaired t-test with Welch’s correction, p<0.01, n=4). This supports the observation of larger neuronal counts per ganglion in the DC. However, to account for differences in neuronal numbers in the PC and DC, the raw neighbor counts were normalized to total neuron count for each region. This revealed significant regional differences and a shift in the neuronal distribution (Fig. 3E). In the DC, a significant proportion of neurons had fewer neighbors, which is indicative of smaller ganglion size and, consequently, smaller neuronal clusters. The larger ganglion size and neuronal clusters were evident from the larger proportion of cells having more neighbors (**Fig. 3E** and **F**) in the PC relative to the DC (normalized counts: 0.27 ± 0.01 vs. 0.23 ± 0.01 for 3 neighbors, 0.14 ± 0.005 vs. 0.2 ± 0.007 for 5 neighbors, 0.05 ± 0.004 vs. 0.1 ± 0.013 for 6 neighbors for DC vs. PC respectively, Mean ± S.D., unpaired t-test with Welch’s correction, p<0.001, p<0.0001 and p<0.01 respectively, n=4). Thus, the number of PNs is proportional to the cell density and size of the ganglion.

Region-specific differences in the distribution of neuronal subtypes could be reflective of the specific functions of the GI subregions (Hamnett, Dershowitz, Sampathkumar, et al. 2022; Li et al. 2019; Nestor-Kalinoski et al. 2022). The regional distribution of the calcium-binding proteins, Calb+ and Calret+ neurons of the PC and DC were examined using images from the Hamnett *et al*. (2022) dataset. The aim was to test the capacity of GAT to detect established regional differences in ENS distribution and investigate the relative distribution of these neuronal markers as they were co-labeled in the same tissue (Fig. 3G). Using GAT analysis, a higher proportion of Calb+ neurons were detected in the PC compared to the DC (Fig. 3H; 1.99 ± 0.78 for DC vs 6.25 ± 2.71 for PC, Mean ± S.D., unpaired t-test with Welch’s correction, p<0.05, n=4). Conversely, no significant differences in the number of Calret+ neurons were detected across regions (Fig. 3I; 26.96 ± 4.9 for DC vs 23.51 ± 9.3 for PC, Mean ± S.D., unpaired t-test with Welch’s correction, p>0.5, n=4) consistent with Hamnett *et al*. (2022). In this dataset, preparations were co-labeled for Calb and Calret, enabling the use of GAT to assess neurons that were double positive for these markers. A very small proportion of Calret and Calb double positive neurons were detected in PC and DC, and no significant difference was found in the distribution across regions (Fig. 3J; 0.65 ± 0.46 for DC vs 1.54 ± 0.71 for PC, Mean ± S.D., unpaired t-test with Welch’s correction, p>0.05, n=4).

The distribution of Calb+ and Calret+ neurons was further assessed using spatial analysis in GAT. This revealed significantly greater numbers of neurons with one Calb+ neighbor in the PC compared to the DC (Fig. 3K; 0.18 ± 0.06 for PC vs. 0.06 ± 0.02 for DC, Mean ± S.D., unpaired t-test with Welch’s correction, p<0.05, n=4). This finding aligns with the observation that the PC contains a greater proportion of Calb+ neurons (Fig. 3H), resulting in a larger number of Calb+ neighbors surrounding each neuron compared to DC. No difference between regions was detected for neurons with Calret+ neighbors (**Fig. 3I**), which aligns with the finding of lower proportion of Calret+ neighbors in both DC and PC. The spatial distribution of neurons that coexpressed Calb+ and Calret+ was also determined, with no significant difference between regions (**Fig. 3M-N**). Spatial analysis of Calret+ neurons relative to Calb+ neurons and vice versa did not reveal any regional differences, suggesting that Calret+ neurons do not preferentially associate with Calb+ neurons in either the PC or DC. These analyses demonstrate that spatial analysis using GAT effectively detects regional differences in neuronal and neuronal subtype distribution.

## Discussion

GAT is a user-friendly Fiji-based software for studying the distribution of enteric neurons and their neurochemical coding in whole mounts of GI tissue in 2D. It uses DL models for segmentation of neurons and ganglia which enables higher throughput and faster data extraction making it possible to analyze large tissue areas. Moreover, spatial analysis in GAT provides an objective means for characterizing the distribution of cells and distinct subpopulations. The DL models are coupled with a user-friendly workflow thus enabling researchers with minimal computational expertise to adopt GAT for rapid and reproducible image analysis.

The availability of state-of-the-art user-friendly tools such as StarDist (Schmidt et al. 2018) and deepImageJ (Gómez-de-Mariscal et al. 2021), in combination with ZeroCostDL4Mic (von Chamier et al. 2021) was crucial for the development of GAT. The GAT enteric neuron models can be used within any software that supports StarDist, thus giving the user flexibility to generate custom analysis pipelines should they be required. The GAT software repository has models and scripts compatible with QuPath, a popular image analysis software for analyzing whole slide images and large 2D images. Cellpose was used to compare the segmentation abilities of the DL models as it works readily on diverse datasets and the user interface makes it simple to use. It is unclear if the cyto2 model was trained on enteric neuronal datasets, which may explain the slightly lower performance metrics when compared to GAT (**Fig. 2**). In Cellpose v2.0, additional models trained on fluorescent images are available, which may show an improvement over cyto2 (Pachitariu & Stringer 2022). Moreover, the best performance can be achieved by fine-tuning the Cellpose models, where the GAT training dataset can be combined with user-specific data to generate custom models (Pachitariu & Stringer 2022). This could increase the accuracy of the segmentation. However, performing quality checks with objective metrics is essential, as shown in **Fig. 2A, 2C**, and **S6**.

StarDist was used for segmenting cells within GAT, as the software is optimized to detect objects with star-convex shapes such as cell nuclei. Thus, it is suitable for detecting enteric neurons that have a circular shape (Schmidt et al. 2018). Other important benefits include: the availability as a Fiji plugin; the ability to use StarDist within macros/scripts; the ability to tune the cell detection by changing the “probability” value, allowing segmentation of images with varying labelling intensities or with high background noise and the ability to detect overlapping cells. The overlap detection can be adjusted by changing the “overlap threshold” value. This is particularly useful for analyzing tissue that has not been stretched appropriately or tissue from larger animals where the ganglia are thicker resulting in greater overlap between cells in 2D. The caveat of using StarDist is that it can only be used for round cells and not for cells with complex morphology. This currently limits its use to enteric neurons, and other celltypes where a nuclear stain is available, such as Sox10 for enteric glia. Thus, other cells with non-circular complex shapes, such as tissue resident macrophages or interstitial cells of Cajal, cannot be analyzed using the current pipeline. Future versions of GAT aim to add support for analyzing diverse cell types within the gut wall.

A limitation of using DL-based models in GAT is that they may not work across image types that GAT has not previously encountered. This could be rectified by retraining the models with new data, but it may not always be feasible as this process is laborious and requires computational expertise. Given the evolving landscape of image analysis and cell segmentation software, GAT offers an option of importing custom segmentations for cells and ganglia directly into the analysis pipeline. This feature allows flexibility for the user to choose their preferred cell segmentation tool. As an example this approach was used in **Fig. 3**. The neuron subtype model successfully segmented Calb+ neurons but was not consistent for Calret+ neurons as the labelling was heterogenous. Similarly, ganglia segmentation was not consistent using the ganglia model. To rectify this, QuPath was used for training an object classifier for Calret+ neurons and a pixel classifier for ganglia, thus enabling segmentation of ganglia and Calret+ neurons respectively. The respective detections and annotations were exported from QuPath and reimported back into GAT during analysis.

One limitation of GAT is that the analysis workflow is currently designed for 2D images, as the Fiji StarDist plugin (v0.3.0) is limited to 2D datasets. Currently, GAT does not support importing 3D segmentation from other software. As cells which occupy a 3D space are projected onto a 2D image, they can often superimpose or overlap with each other, leading to challenges in accurately delineating and segmenting these cells. Separation of cells is more readily achieved with 3D data; however, only the Python implementation of StarDist supports 3D segmentation. Furthermore, cell shape and size are more accurately measured in 3D compared to 2D. One reason is that the volume measurements are less impacted by differences in tissue stretch compared to area measurements in 2D. 3D segmentation requires DL-based approaches that use high-quality 3D annotations, which is a time-consuming and laborious process. Several tools are available for annotating data in 3D (Berg et al. 2019; Boergens et al. 2017; Borland et al. 2021; Fedorov et al. 2012; Tasnadi et al. 2020). However, the success of DL models relies on the availability of large amounts of high-quality training data, which account for the diversity of the markers used, animal species studied, how the tissue is prepared, and the variability in the instruments used for acquisition. Existing tools such as Cellpose (Stringer et al. 2021), 3D ImageJ suite (Ollion et al. 2013), and CLIJ or pyclesperanto (Haase, Jain, et al. 2020; Haase et al. 2022; Haase, Royer, et al. 2020) can be used for processing 3D data and enabling curation of labeled data. To enable ENS-specific analytical solutions, it is essential to have robust training data made accessible to the wider research community. For example, the GAT training dataset was deposited on Zenodo, an open data repository. This has been utilized to generate custom cell segmentation models for image analysis software to quantify Ca^2+^ signaling in the gut (Barth et al. 2023).

Manual analytical approaches to assess changes in the ENS are limited to cellular density and the number of neuron subtypes within the tissue. However, incorporating spatial analysis can reveal insights into cellular distribution, interactions, and potential implications for function at a tissue-level (Nestor-Kalinoski et al. 2022). Existing spatial analysis software solutions require significant computational expertise and may require optimization to study the ENS (Feng et al. 2023; Rose et al. 2020; Stoltzfus et al. 2020; Yang et al. 2020). Factors such as operator expertise, animal species being studied, and the pathology investigated can affect how the tissue is stretched and prepared as a wholemount (Gomez-Frittelli, Hamnett & Kaltschmidt 2023; Kapur 2013). This, in turn, can affect analysis and interpretation of cellular distribution (**Fig. 3A-C**). GAT accounts for possible variations in tissue stretch by using an enteric neuron-specific nearest neighbor distance threshold value. Differences in neuronal distribution were demonstrated objectively using the proximal neighbor analysis in GAT. The PC had relatively larger neuronal clusters compared to the DC indicative of the large and small average ganglia sizes, respectively. Altered distribution of Calret+ and Calb+ neurons across the PC and DC were reflected in the spatial analysis, and no spatial association was found between neuronal subtypes. This approach can be extended to assess changes in the structure or morphology of the ENS across gut regions or in diseases such as inflammatory bowel disease (Brierley & Linden 2014), diabetes (Chandrasekharan et al. 2011; Ingrid et al. 2013) and enteric neuropathies, such as Hirschsprung disease (Heuckeroth 2018). For example, this analysis could be used to determine if there are subtle differences, such as fewer nNOS+ neuronal neighbors, which would be indicative of lower nNOS+ neurons in these conditions. The parameters, such as the average PN distance, may need to be modified for human tissue, and the option to test different values is available in GAT. Due to the analysis capabilities of GAT being limited to enteric neurons, the spatial analysis does not capture the complexity of cellular distribution and relationships with other cell types, such as enteric glia, macrophages, and interstitial cells of Cajal. Incorporating additional spatial analysis metrics such as those related to cell colocalization, spatial heterogeneity (Feng et al. 2023), or the use of a spatial neighbors graph to understand neighborhood enrichment (Palla et al. 2022) could enable a more comprehensive and unbiased examination of cellular interactions and distributions in the gut.

GAT excels at segmenting neurons and works across images of varying staining qualities. It offers a faster alternative to manual analysis and has been designed with ENS-specific analysis solutions, such as studying neurochemical coding and spatial analysis of neuronal distribution. New features are regularly being introduced based on user feedback and experience. To enable compatibility for further analysis in other software, segmentation maps, cell outlines, cell type information, and cell coordinates are extracted for each experiment during analysis. GAT is written in Fiji using the macro language, limiting the scope of the user interface (UI) and its interactivity. Future versions of GAT may use scripting languages in Fiji, which offers more flexibility in developing highly interactive UIs. Moreover, the availability of StarDist models enables the development of interactive workflows in napari and QuPath. GAT aims to create a toolbox that automates common image analysis tasks in ENS research, eliminating the burden of manual analysis. This allows scientists to spend more time interpreting biology and advancing scientific research.

## Methods

### Mice

The details on housing conditions and ethics for the mice used within this study can be found in the respective studies (DiCello et al. 2020; Hamnett, Dershowitz, Sampathkumar, et al. 2022; McQuade et al. 2021; Nestor-Kalinoski et al. 2022; Poon et al. 2022).

B6;129S6-*Gt(ROSA)26Sor^tm9(CAG-tdTomato)Hze^*/J (Jackson Laboratory, Bar Harbor, ME; stock no. 007905) mice were crossbred with B6.Cg-*E2f1^Tg(Wnt1-cre)2Sor^*/J (Jackson Laboratory; stock no. 022501) mice. The F1 offspring expressed tdTomato within all ENS cells. The animals were handled in accordance with the institutional guidelines of the University of Tübingen, which conform to international guidelines. Mice were housed in standard plastic cages with standard bedding under a 12-hour light-to-dark cycle at 22 ± 2°C, 60% ± 5% humidity, with free access to food and water.

### Rat

The details on housing conditions and ethics for the rats can be found in (Furness et al. 2023).

### Human Tissue

Pediatric colon tissue was obtained from 1 patient, (male, 4months old with rectosigmoid disease with prior written informed consent (Royal Children’s Hospital and Monash University (HREC: 38262 and 21091)). The ethics for the adult human tissue can be found in Chen et al. (2023).

### Whole mount tissue preparation and immunohistochemistry

#### Mouse

The protocols used for dissection and immunohistochemistry of mouse myenteric wholemounts varied across labs and are summarized in the respective publications from each source (DiCello et al. 2020; Hamnett, Dershowitz, Sampathkumar, et al. 2022; McQuade et al. 2021; Nestor-Kalinoski et al. 2022; Poon et al. 2022).

#### Human

The protocol for human colon samples can be found in Chen *et al*. (2023).

**Table 1.**
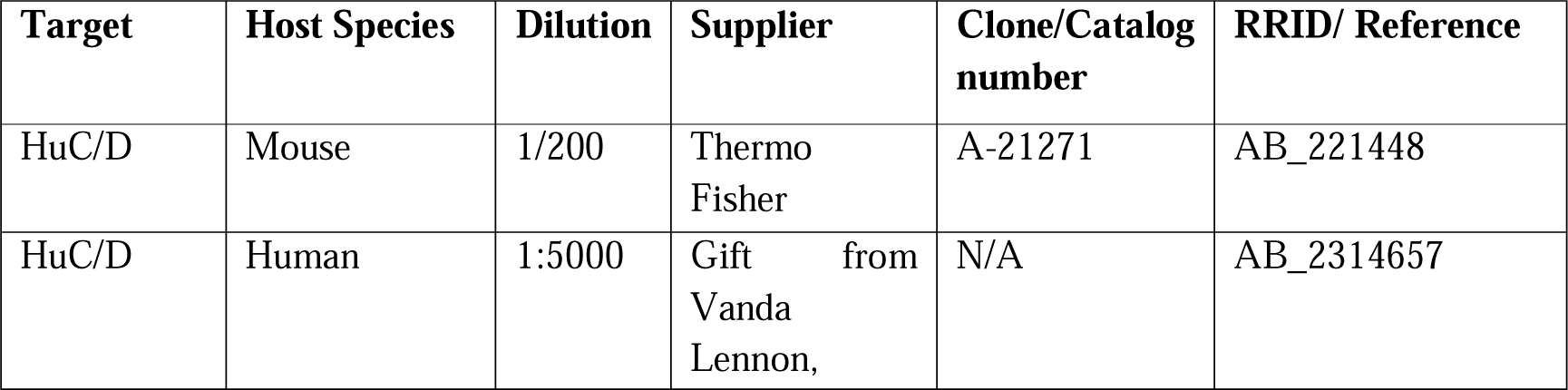
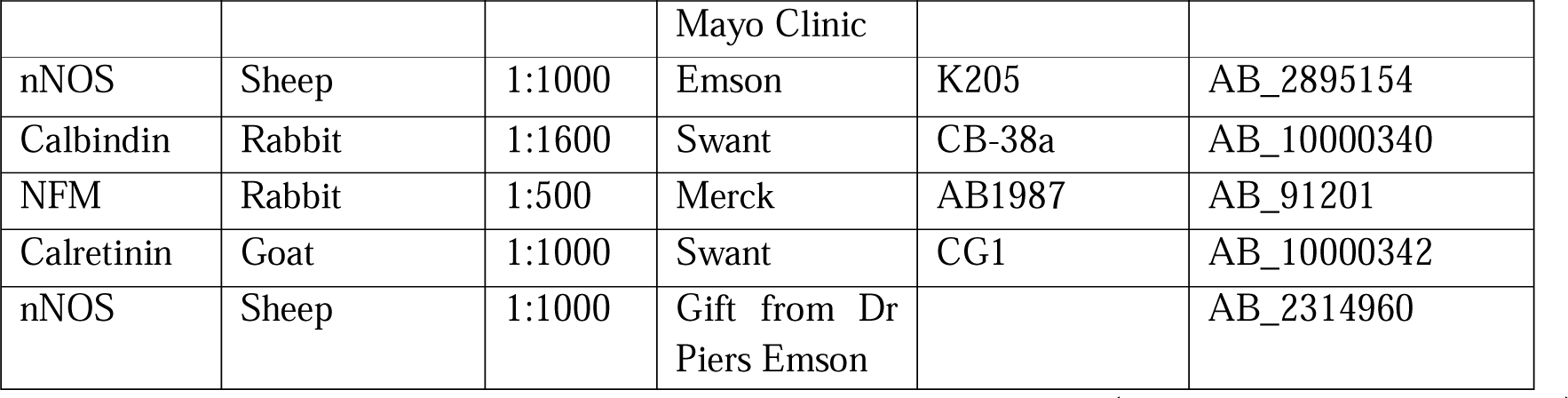
Primary antibodies used for immunofluorescent labeling of wholemount preparations.

| Target | Host Species | Dilution | Supplier | Clone/Catalog number | RRID/ Reference |
| --- | --- | --- | --- | --- | --- |
| HuC/D | Mouse | 1/200 | Thermo Fisher | A-21271 | AB_221448 |
| HuC/D | Human | 1:5000 | Gift from Vanda Lennon, | N/A | AB_2314657 |
|  |  |  | Mayo Clinic |  |  |
| nNOS | Sheep | 1:1000 | Emson | K205 | AB_2895154 |
| Calbindin | Rabbit | 1:1600 | Swant | CB-38a | AB_10000340 |
| NFM | Rabbit | 1:500 | Merck | AB1987 | AB_91201 |
| Calretinin | Goat | 1:1000 | Swant | CG1 | AB_10000342 |
| nNOS | Sheep | 1:1000 | Gift from Dr Piers Emson |  | AB_2314960 |

The secondary antibodies used were donkey anti-human 594 (Jackson ImmunoResearch), donkey anti-sheep 647 and donkey anti-sheep 488 (Thermo Fisher Scientific). Streptavidin-AMCA was used with donkey anti-mouse biotin for Hu in the adult human colon preparations (Chen et al. 2023).

#### External Datasets for model training

External datasets were also used for creating DL models. Data from the Stimulating Peripheral Activity to Relieve Conditions (SPARC) program website (https://sparc.science) were used for training the neuronal, neuronal subset and ganglia segmentation models. Within the SPARC portal, data from mouse (Nestor-Kalinoski et al. 2022; Wang et al. 2021) and human myenteric plexus whole mounts (Chen et al. 2023; Graham et al. 2020b) were used for curating training datasets.

#### Image acquisition

Datasets generated for this study were acquired using different instruments. This enables the DL models to generalize well and work across a variety of data acquired with commonly used microscopes.

The images for the mouse tissue were acquired using:

- Leica TCS-SP8 confocal system (20x HC PL APO NA 1.33, 40 x HC PL APO NA 1.3)
- Leica TCS-SP8 Lightning confocal system (20x HC PL APO NA 0.88)
- Zeiss Axio Imager M2 (20X HC PL APO NA 0.3)
- Zeiss Axio Imager Z1 (10X HC PL APO NA 0.45)

Human tissue images were acquired using:

- IX71 Olympus microscope (10X HC PL APO NA 0.3)(Chen et al. 2023)
- Leica TCS-SP8 confocal system (20x HC PL APO NA 1.33, 40 x HC PL APO NA 1.3)

Acquisition conditions for SPARC datasets are:

- Leica TCS SP5 laser scanning confocal microscope (20x NA 0.70, 40x NA 1.25, or 63x NA 1.4) (Nestor-Kalinoski et al. 2022)
- Zeiss LSM 710 confocal microscope (10x and 20x PL APO). Z-axis increments were 4μm (10x objective) or 1μm (20x objective) (Graham et al. 2020b).

### Software

#### Training data

Training data were generated using custom scripts written in Fiji macro language, provided with GAT (GAT -> Tools -> Data Curation). Briefly, the entire image or a portion of the image was selected for annotation. This was followed by manually outlining neurons using the drawing tools and saving them in the ROI Manager. For the enteric neuron datasets, the annotated images of varying sizes were saved as raw images and as segmented label images, where each neuron had an individual pixel value. In the label images all pixels with value 1 belong to neuron 1, all pixels with value 2 belong to neuron 2 and so on. The binary masks were saved for ganglia segmentation. The annotation was performed and verified by at least two researchers experienced with the identification of enteric neurons and ganglia. Briefly, the outlines from the label image were overlaid on the raw images and verified for each cell inusing Fiji. The “Verify Images Masks” macro within GAT -> Tools -> Data Curation were also used. For images that had DAPI labelling, the neuronal nuclei were used to delineate overlapping cells. Incorrect ROIs were deleted and redrawn using the Oval or Freehand drawing tools in Fiji.

#### Enteric neuron models

The training images (**Tables S1 and S2**) were normalized to account for the any variations in the sizes of cells in pixels due to image acquisition conditions, such as resolution and magnification, and species differences. The images from mouse and rat tissue were scaled to a pixel size of 0.568 µm per pixel, whereas the images from human tissue were scaled to 0.9 µm per pixel due to the larger cell sizes. This rescaling process ensured that the training images had a uniform average neuron area of 701.2 ± 195.9 pixel^2^ (Mean ± SD, 6267 cells) irrespective of image magnification or animal species. A similar approach was used to generate a training dataset for the enteric neuron subtype model. This training dataset contained images of neurons expressing various neurochemical markers, including Calb, nNOS, CalR, choline acetyltransferase (ChAT), delta-opioid receptor (DOR), mu-opioid receptor (MOR), neurofilament 200 (NF200) and somatostatin. The average cell area in the neuronal subtype dataset was 880.9 ± 316 pixel^2^ (Mean ± SD, 924 cells), with around 56.6% of the cells being nNOS positive. Thus, nNOS cells were overrepresented in the dataset. The StarDist v0.3.0 Fiji plugin was used for inference.

#### Ganglia model

The ganglia model was trained on images (**Table S3**) with both the pan-neuronal marker, Hu and a second marker that labeled the neuronal or glial fibers. Regions where both markers were co-distributed were manually labeled as ganglia. The markers used for the identification of ganglionic structures consisted of any of the following: Protein Gene Product 9.5 (PGP9.5), nNOS, glial fibrillary acid protein (GFAP), S100b, Tuj1, or NF200. The DeepImageJ v2.1.12 Fiji plugin was used for inference.

#### Deep learning models and software

StarDist v0.7.3 (Schmidt et al. 2018) was used via ZeroCostDL4Mic v1.13 notebooks (von Chamier et al. 2021) within Google Colab for training the 2D segmentation models for enteric neurons and neuronal subsets. The ganglia model was trained in Google Colab using a 2D UNet architecture (Ronneberger, Fischer & Brox 2015) and exported to be readily used within deepImageJ (Gómez-de-Mariscal et al. 2021). The notebooks used for training the models, the training datasets used, training reports, model quality reports, and the models are deposited online at: https://doi.org/10.5281/zenodo.6096664.

Cellpose (v0.7) has been used as a baseline for comparing cell segmentation in this study. It is a generalist cell segmentation solution aimed at analyzing a wide variety of cell types (Stringer et al. 2021). It is not known if Cellpose has been trained on enteric neuron images.

All training data and deep learning models are deposited on Zenodo: https://zenodo.org/doi/10.5281/zenodo.6094887.

#### COUNTEN analysis

For benchmarking with COUNTEN (Kobayashi et al. 2021), the software was accessed from: https://github.com/KLab-JHU/COUNTEN. The analysis used the default values of sigma and the minimum number of neurons per ganglia, set at 7 and 3 respectively. An Google Colab notebook was designed to enable interactive analysis with an option for batch analysis, which can be accessed from: https://github.com/pr4deepr/COUNTEN.

#### Analysis of calbindin and calretinin positive cells

A publicly available dataset (Hamnett, Dershowitz, Gomez-Frittelli, et al. 2022) was used for analyzing Calb and CalR-positive cells. Image files with Calb and CalR co-labelling (EXP 174) were analyzed using a combination of QuPath v0.4.3 (Bankhead et al. 2017) and GAT. Due to inconsistent segmentation of the ganglia using the ganglia model, a pixel classifier was trained in QuPath to identify ganglia based on the co-expression of Hu, Calb and CalR. The resulting annotations were exported from QuPath as Fiji-compatible ROIs, which were subsequently imported into GAT for analysis. Similarly, the enteric neuron subtype model detected CalB-positive neurons but did not reliably detect CalR-positive cells. To address this, an object classifier was trained in QuPath to detect CalR-positive neurons. The neurons were initially detected using the enteric neuron StarDist model based on the Hu channel, followed by the application of the object classifier. Only the calretinin-positive ROIs were extracted from QuPath and imported into GAT for analysis.

The results for each replicate were merged into summary data using the scripts within GAT - > Analysis. The summary data were analyzed in Python (v 3.9.15) and pandas (v 2.0.2) and visualized using seaborn (v 0.12.2). The analyzed data were exported, and statistical analysis was performed in GraphPad Prism (v 9.5.1).

#### Other

ChatGPT (GPT-3.5) was used for initial formatting and editing of the manuscript.

## Supplementary

**Figure S1:**
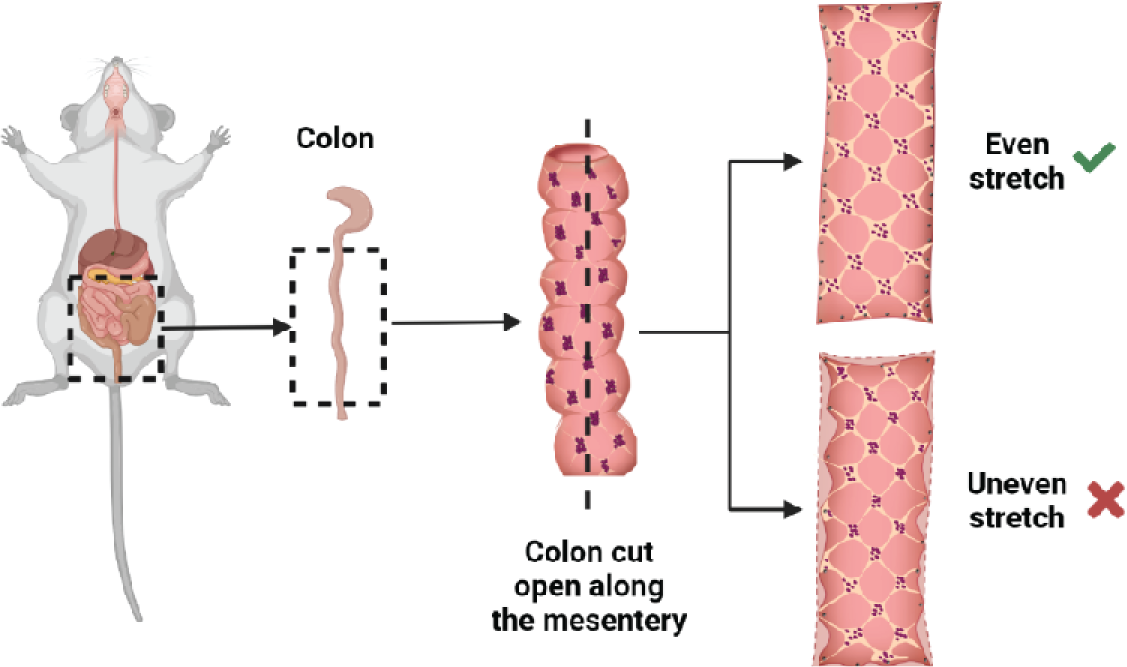
Effects of sample preparation and tissue stretch on the ENS. For wholemount preparations, the isolated intestine is cut open along the mesenteric border, stretched, and pinned as a flat sheet. Individual expertise can determine the quality of this dissection and preparation (Some parts of the image were created with Biorender.com).

**Figure S2.**
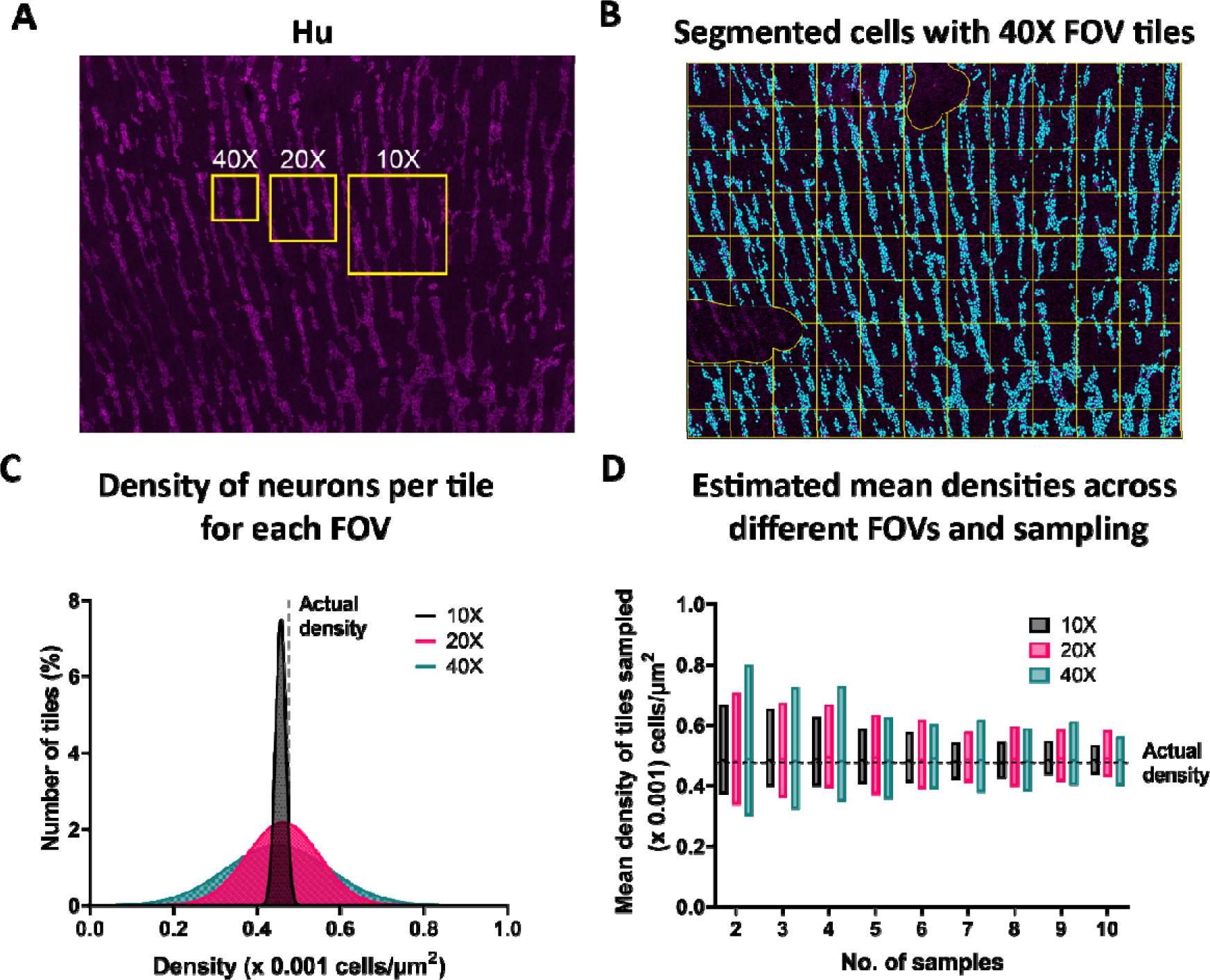
Neuronal density estimates are highly variable at lower magnifications and sampling. **(A)** An image of mouse colon tissue labeled for the pan-neuronal marker Hu (cell density 13.9 mm^2^ area). Overlays represent the area occupied by different magnifications (10X, 20X, and 40X). **(B)** Using a StarDist-trained model, QuPath was used to segment 6650 neurons (blue overlay). Regions with inconsistent staining or dissection were excluded (yellow overlay). Each yellow tile is representative of a 40X field of view (FOV) (**C)** For each individual tile, the density of neurons was calculated to demonstrate that 20X and 40X had a larger variation compared to 10X. **(D)** To illustrate the effect of sampling on averaged neuronal density estimates, images from each magnification were randomly sampled and mean cell densities were estimated across different sampling rates (2-10 images). This was repeated 100 times for each FOV and sampling number. Each bar in the graph represents mean estimates of mean densities for each sampling and FOV choice. Bars represent the minimum and maximum value. The black dotted line indicates the cell density calculated from the entire field of view in **A**, 0.4768 (x 0.001 cells/µm^2^).

**Figure S3:**
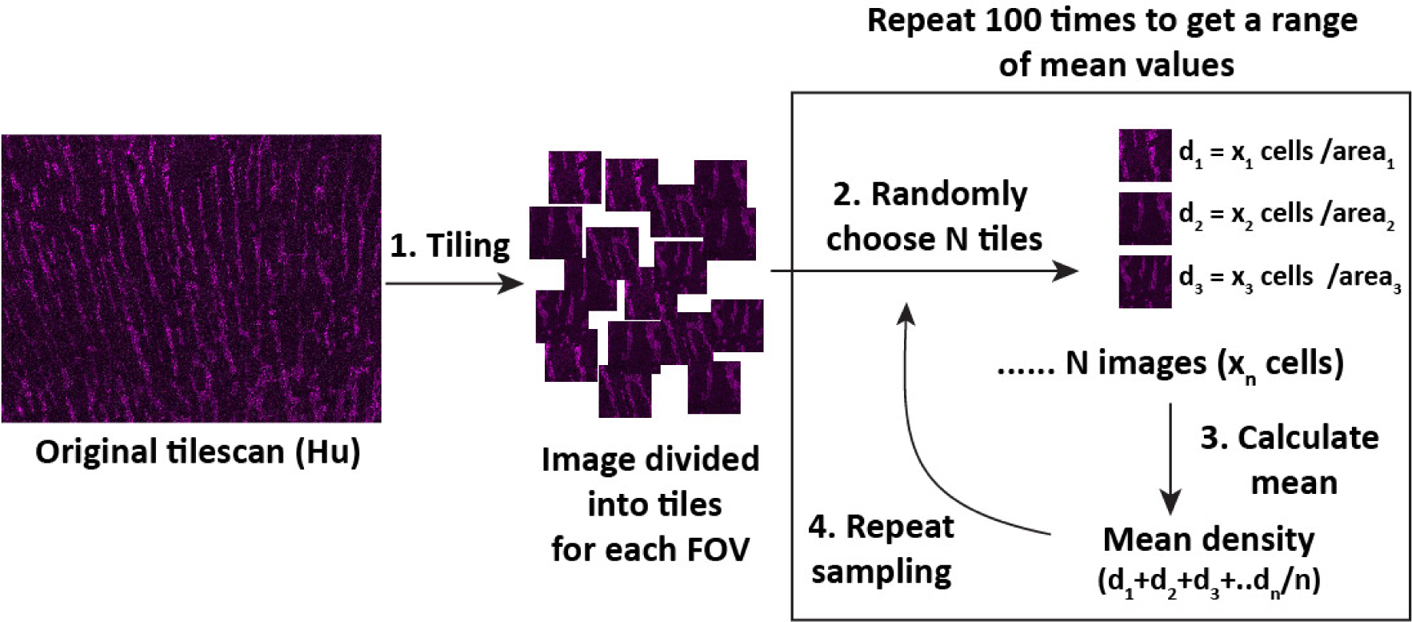
Random sampling of tiles to determine mean cell counts for different FOVs. An illustration for **Fig. S2D** outlining how calculations were performed. The image with Hu labelling was divided into tiles for each FOV (10X, 20X, or 40X). To simulate experiments for a specific magnification with different sample numbers, a range of samples from 2 to 10 was chosen. The mean cell density was calculated for each sample number, and this was repeated 100 times to estimate the range of mean values that could be obtained.

**Figure S4:**
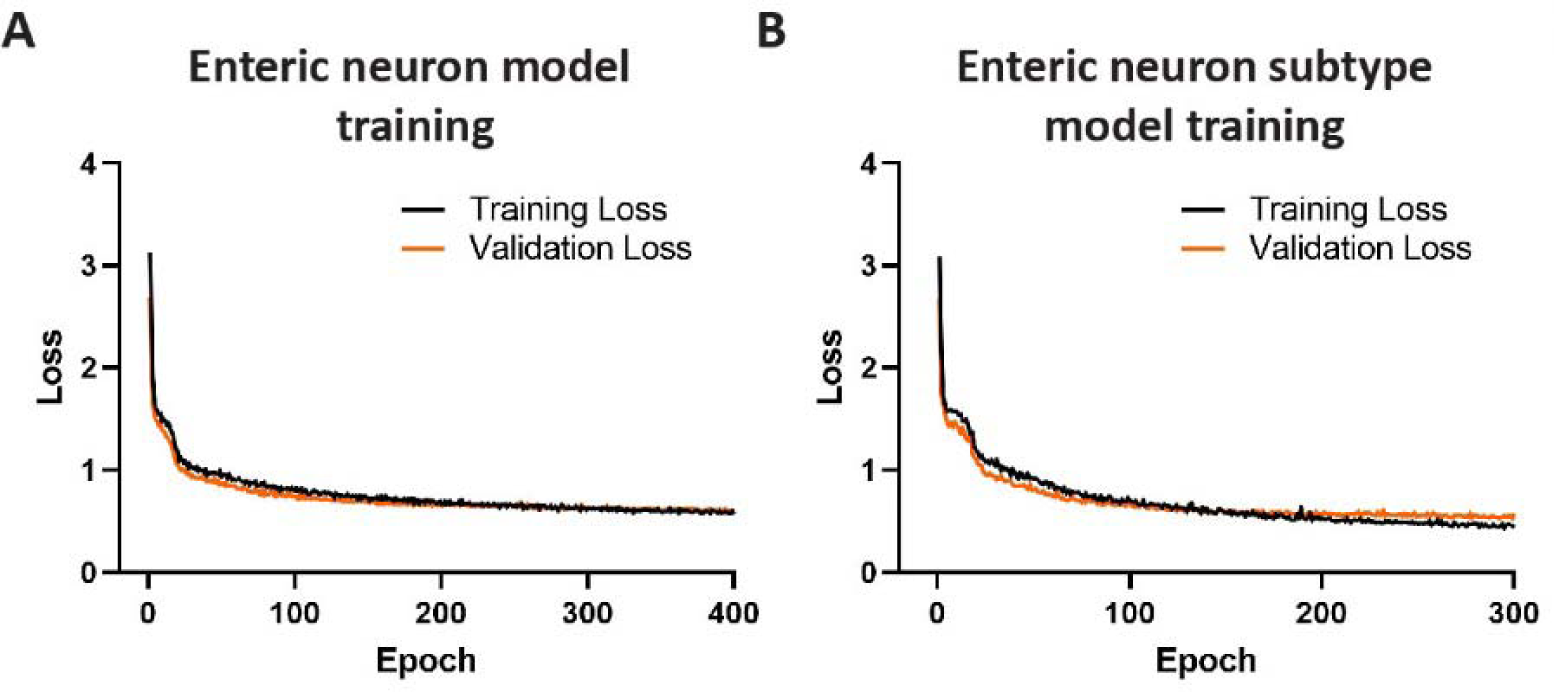
Training and validation loss for different models. Training loss of the StarDist enteric neuron model (A) and enteric neuron subtype model (B).

**Figure S5:**
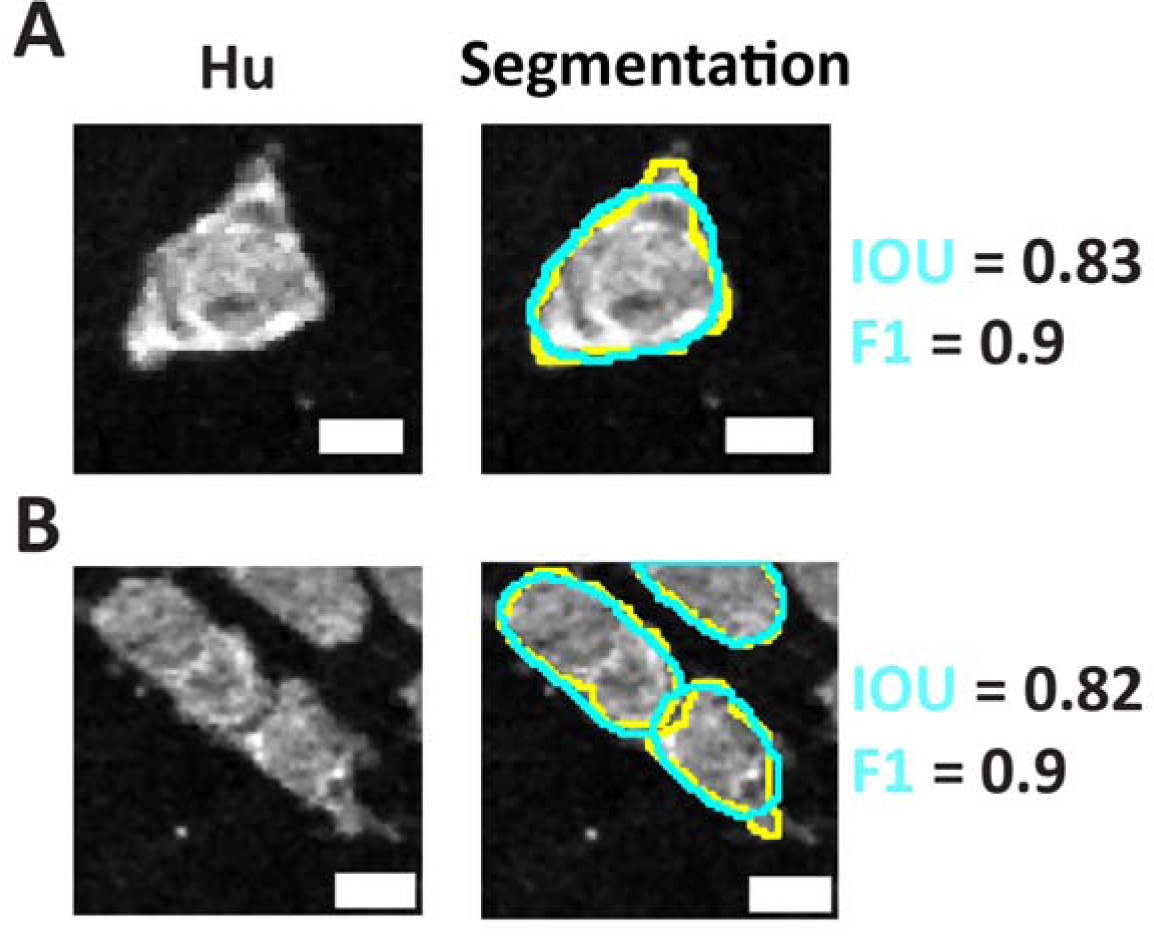
Comparison of StarDist segmentations. Comparison between representative StarDist (cyan outline/text) segmentations and manual annotation (yellow outline). The StarDist predictions have smoother boundaries, but do not capture the original cell contours (Scale bar = 10 µm).

**Figure S6:**
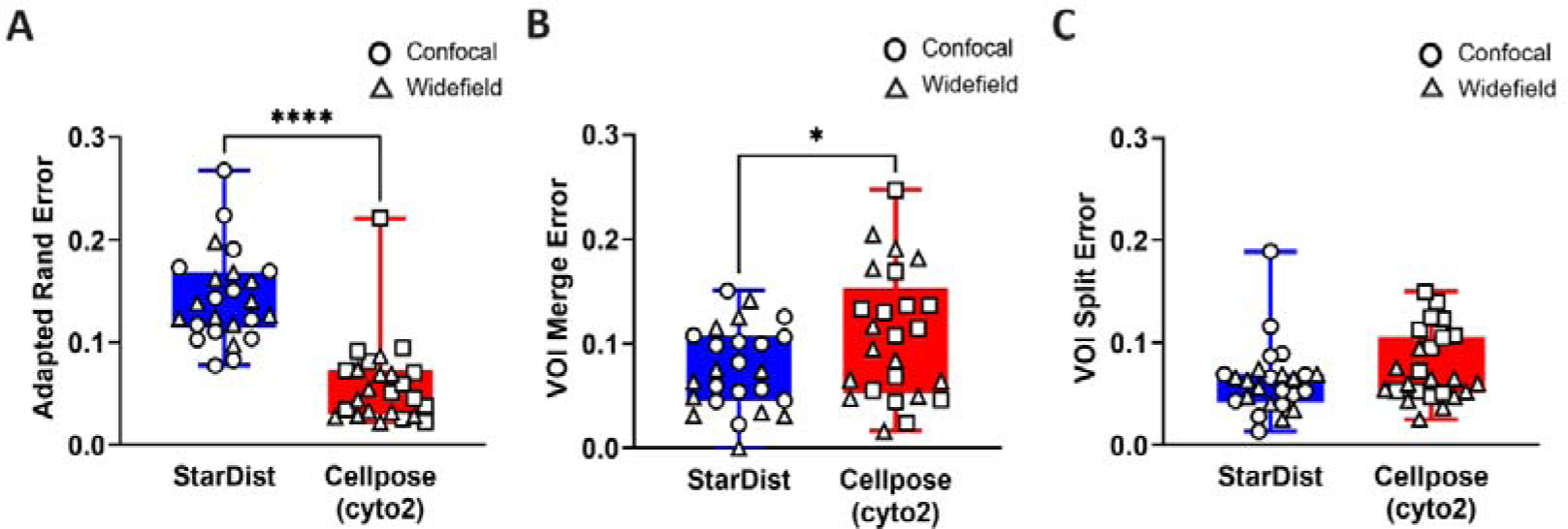
Evaluation of Cellpose and StarDist segmentation metrics for enteric neurons. Cellpose had a lower Adapted Rand Error indicating higher overall segmentation quality than the StarDist model (**A**). However, this could be because Cellpose is better at predicting accurate cell borders compared to how StarDist approximates the cell shape using star convex polygons. The Cellpose model had larger merge and split errors (**B** & **C**) which may explain the higher percentage cell count error presented in **Figure 2**.

**Figure S7:**
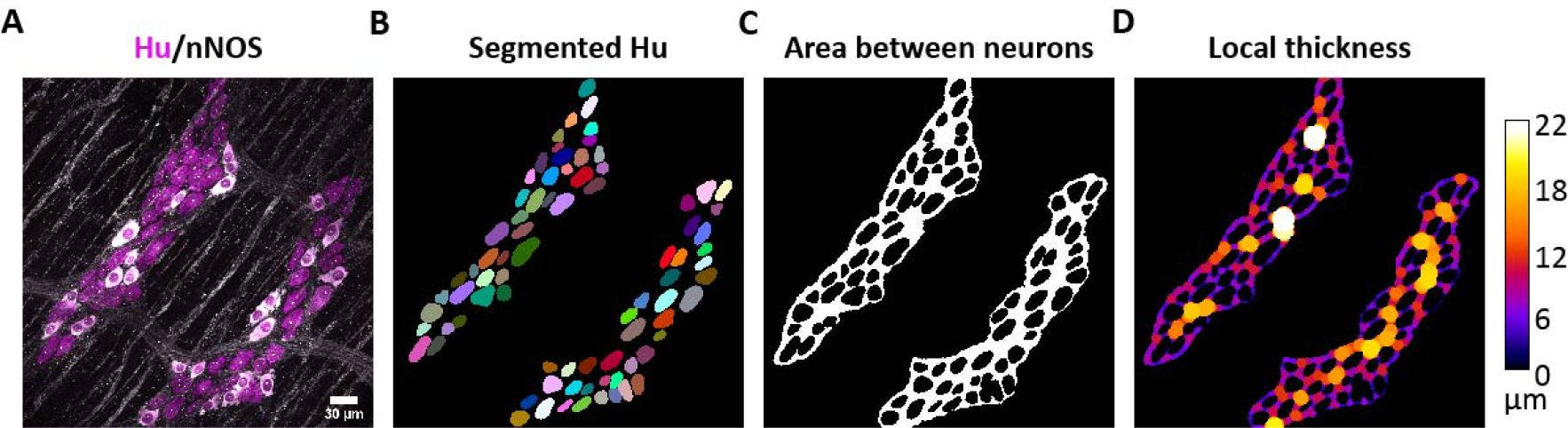
Average proximal neighbor distances can be determined by calculating the local thickness within the ganglia. The distances between neurons in a ganglion can be determined by segmenting the neurons and the ganglia (**A, B**) and measuring the ganglionic area, excluding neuronal soma **(C)**. **(D)** The local thickness map can be used to derive the interneuronal distances. This was calculated by applying the “Local Thickness” plugin (Dougherty & Kunzelmann 2007) on the binary image **(C)**. The “Local Thickness (complete process)” option in Fiji was used to compute the local thickness map as shown in **D**.

**Table S1:** Summary of the images used for training the enteric neuron StarDist model. All images contain cells labeled with the pan-neuronal maker Hu. The number of images are mentioned under the species and Total images.

| Species | Lab 1 | Lab 2 | Lab 3 | SPARC | Lab 4 | Total images |
| --- | --- | --- | --- | --- | --- | --- |
| Mouse | 44 | 15 | 6 | 1 | - | 66 |
| Rat | 2 | - | - | - | - | 2 |
| Human | 2 | - | - | 6 | 8 | 16 |
| Magnification | 40X/20X | 10X | 20X | 20X/40X | 10X | - |
| Modality | Confocal | Widefield | Widefield | Confocal | Widefield | - |
| Total images | 48 | 15 | 6 | 8 | 7 | 84 |

**Table S2:** Number of images used for training the enteric neuron subtype StarDist model.

| Data source | nNOS | MOR | DOR | ChAT | CalR | CalR | NFM | CGRP | SST | Total/<br>source |
| --- | --- | --- | --- | --- | --- | --- | --- | --- | --- | --- |
| Lab 1 | 4 | 2 | 1 | 2 | 1 | - | - | - | - | 10 |
| Lab 2 | 16 | - | - | - | - | - | - | - | - | 16 |
| Lab 3 | 3 | - | - | - | 5 | 9 | 6 | - | - | 23 |
| SPARC | 1 | - | - | 3 | - | 1 | - | 1 | 1 | 7 |
| Total/<br>marker | 24 | 2 | 1 | 5 | 6 | 10 | 6 | 1 | 1 | 56 |

**Table S3:** Number of images used for training the ganglia model. Each marker was used in combination with Hu to generate training data for the 2D U-Net ganglia model implemented using deepImageJ.

| Data source | PGP9.5 | GFAP | Wnt1Cre-GFP | ChAT | NF200 | 5HT | CGRP | NPY | nNOS | s100b | Tuj1 | Total / source |
| --- | --- | --- | --- | --- | --- | --- | --- | --- | --- | --- | --- | --- |
| Lab 1 | 12 | 28 | 3 | - | - | - | - | - | 5 | - | - | 48 |
| Lab 2 | - | - | - | - | - | - | - | - | 12 | - | - | 12 |
| Lab 3 | - | - | - | - | - | - | - | - | - | - | - | 0 |
| Lab 4 | - | - | - | 1 | 1 | 2 | 1 | 1 | - | - | - | 6 |
| SPARC | - | - | - | 4 | - | - | - | - | - | 3 | 3 | 10 |
| Total/ marker | 12 | 28 | 3 | 5 | 1 | 2 | 1 | 1 | 17 | 3 | 3 | 76 |

## Supplementary methods

### Effects of varying magnification and sampling

An image of a myenteric wholemount of the mouse colon (13.9 mm^2^) labeled with Hu was used with QuPath v0.3.2 (Bankhead et al. 2017) to test the effects of varying magnification and sampling on estimated cell counts (**Figs 2** & **3**). A whole image annotation was created and then divided into tiles, where each tile had areas of 775,918 µm^2^, 338,116 µm^2^ and 150,274 µm^2^, thus simulating 10X, 20X, and 40X magnifications, respectively. Tiles at the edges of the tissue that were below 60,000 µm^2^ in area and areas with uneven staining were excluded in the subsequent calculations. To perform cell segmentation, a custom groovy script in combination with the StarDist 2D enteric neuron model converted into ONNX format was used in QuPath (https://github.com/pr4deepr/GutAnalysisToolbox/tree/main/QuPath_workflow). The parameters used for segmentation were a rescaling factor of 1 and a probability of 0.65. Once segmentation was performed, the cell numbers were saved using the annotation measurement tables. Once cell counts were estimated, a custom Python script was used to choose random tiles and estimate mean values. This is illustrated in **Figure S1**. The Python code is provided as a Jupyter notebook.

### Evaluation of CellPose and StarDist segmentation

The segmentation metrics for Cellpose cyto2 and StarDist enteric neuron model were further evaluated using Adapted Rand error, VOI merge and VOI split on the test data from the enteric neuron model training. Adapted Rand error assesses the overall segmentation quality, whereas VOI merge and VOI split assess errors associated with cell merging and splitting respectively (Wolny et al. 2020). The evaluation script from plant-seg-tools GitHub repository (https://github.com/hci-unihd/plant-seg-tools/tree/main) was used to evaluate the segmentation results from the Cellpose cyto2 and enteric neuron StarDist model against the ground truth data **(Fig. S6)**.

## Supplementary text

### Effects of sample preparation and tissue stretch on neuronal counts

Gastrointestinal wholemount preparations are typically used to quantify enteric neuron numbers. More accurate estimates can be achieved with these preparations in comparison to tissue sections, as all neurons within the same plane as the plexuses can be visualized within the 3D or 2D space, and larger tissue samples can be prepared to increase sampling. To prepare a wholemount, the tissue is cut open along the mesentery, stretched and pinned for further processing (illustrated in **Fig. S1A)**. This flat sheet preparation allows investigation of the anatomical features across larger tissue areas. Depending on the expertise of the researcher, inconsistent stretching and uneven sampling can lead to biased estimation of cell numbers (**Fig. 3A**). One way to mitigate this effect of stretching is to compare the dimensions of the tissue before and after stretching and include a correction factor for changes in tissue dimensions caused by stretching and any age-associated changes (Gamage et al. 2013). However, very few studies have used this approach (Gamage et al. 2013; Kapur 2013; Michel et al. 2022b).

### Effects of inadequate sampling and image acquisition

Advances in imaging technologies and software have enabled the sampling of large tissue areas as tile scan images, a standard feature in many conventional widefield and confocal microscopes. However, the analysis throughput of large images remains a significant limiting factor.

Due to the time-consuming nature of the manual analysis, the quantification of neurons along larger lengths of the intestine is not standard practice within the field (Michel et al. 2022b; Nestor-Kalinoski et al. 2022). Estimates of neuron densities are calculated from randomly chosen FOVs selected from small sections of the gut, which are averaged. These values are representative of total numbers across entire regions of the intestine. The following section aims to quantify the errors associated with FOV selection and choice of magnification during image acquisition.

A large mouse proximal colon myenteric wholemount (13.9 mm^2^) was labeled for Hu (**Fig. S2A**). GAT StarDist model converted to ONNX format for QuPath was used to calculate a cell density of 0.4768 (x 0.001 cells/µm^2^) across the tissue. To simulate the effects of sampling using single images at 10X, 20X, and 40X, the whole image was divided into smaller tiles equating to these magnifications on a Leica SP8 confocal microscope (**Figs. S2A** & **B)**. The cell count was estimated for each tile and plotted as a frequency histogram (**Fig. S2C**). The density values for 20X and 40X FOVs had a larger variability compared to 10X (Range (max – min): 0.3844 for 10X, 0.5766 for 20X and 0.5913 for 40X). Thus, there is increased variability when estimating cell density with higher magnifications, as only smaller tissue areas can be sampled.

The choice of magnification and sample number could affect the accuracy of the final cell counts. Cell counts were averaged from 2 to 10 tiles randomly selected at each magnification to measure the effects of sample size and subsequently averaged (**Fig. S2C)**. To estimate the range of mean values that could be obtained, this process was repeated 100 times (**Fig. S3**). Each bar in **Fig. S2D** shows 100 possible mean density estimates for all the different sample numbers. The larger the bar, the higher the variability in the final estimate. The actual cell density is calculated for the entire FOV and is denoted by the black dotted line. When comparing 10X and 40X FOVs, 10 images of the 40X FOV were required to have a comparable variability to 5 images from 10X FOV (Range of 0.182 for 10X and 0.27 for 40X for 5 samples vs Range of 0.1 for 10X and 0.164 for 40X for 10 samples). Thus, it is recommended to use larger FOVs, such as a 10X or 20X magnification and increased tissue sampling to ensure accurate cell density estimates.

### Extensibility of GAT models

The StarDist models are compatible with other image analysis software, including QuPath (Bankhead et al. 2017) and napari (Sofroniew et al. 2022). This compatibility enables users to create custom analysis pipelines without being restricted to a single software platform. Some examples include:

- **QuPath** is a popular software for analyzing large 2D images that has an extension for implementing StarDist cell segmentation (Schmidt et al. 2018). The 2D enteric neuron models have been converted into the ONNX format using tf2onnx software (https://github.com/onnx/tensorflow-onnx) to make it accessible from QuPath. A groovy script and the QuPath model are provided in the GAT repository. An advantage of QuPath is its ability to navigate large 2D images easily and use less computing power than Fiji when performing StarDist segmentation on large tile scans.
- **napari** is a popular Python-based multi-dimensional viewer. The availability of an interactive napari StarDist plugin (https://github.com/stardist/stardist-napari) enables deployment of the StarDist enteric neuron models within a Python image processing workflow. Moreover, as the ganglia model is a TensorFlow model, it can be implemented in Python along with the StarDist models.
- **APEER** is a cloud-based image processing platform by Zeiss that allows custom image processing workflows to be run directly from the image acquisition software Zen. It has a StarDist module (https://www.apeer.com/app/modules/StarDist-2D-Nucleus-Segmentation-%28beta%29/52cc57d2-94ea-4417-a4c9-d8d957a3cf32). The GAT StarDist models can likely be used within Zen. Namely, GAT workflows could be set up to perform cell counts immediately after image acquisition.
- **CellProfiler** is a popular cell image analysis software that does not require prior programming experience (McQuin et al. 2018). It currently has a StarDist plugin (https://github.com/CellProfiler/CellProfiler-plugins), which could enable the use of the GAT models within a CellProfiler pipeline.

### Limitation of the DL models

The data used for training DL models is crucial for ensuring the model works accurately across a wide range of images. The DL models are only as good as the training data used, and any biases within the training data are carried over within the model (Jacquemet 2021, Litjens, 2017 #72). For example, earlier versions of GAT models trained on 2D maximum projections of confocal images were not accurate on widefield images. Thus, emphasis was placed on gaining diverse training datasets. This does not necessarily guarantee the robustness of the DL models. Instances where DL models may not work well are discussed below.

### Effect of tissue type, preparation, and species

The training datasets for the models are predominantly from myenteric wholemounts of the mouse colon. As such, the models may not work well on wholemounts of the submucosa, different regions of the gut, or preparations from other species. The GAT enteric neuron models have been tested successfully (not shown) on submucosal regions of the colon, although there are no training data from other layers.

### Effect of imaging modality and choice of fluorophores

As images from a confocal microscope have different noise levels, contrast, and background labeling compared to widefield microscopy images (Shaw 2006), images acquired with both modalities were included in the training dataset. The training data has 60% confocal 2D image projections and 40% widefield images. Moreover, the images consisted of Hu labeling using a range of fluorophores (405, 488, 568, and 647 nm emission). In addition to sample preparation and imaging modality affecting autofluorescence, biological tissue is increasingly autofluorescent at lower wavelengths. Autofluorescence from blood vessels can affect cell segmentation, leading to erroneous segmentation and predictions.

### Neuronal subtype model

The distribution of labeling in the neuronal soma varies between markers in the training dataset used. This includes punctate staining (delta or mu-opioid receptors), cytoplasmic staining (nNOS), or variable staining intensities (CalR, nNOS). Due to the high variability, the neuronal subtype model may not work across all neuronal markers. This issue was encountered for the analysis in **Fig 3**. Increasing the size of the dataset with images for different markers or using a different type of neural network architecture may increase the accuracy of the neuronal subtype predictions. Alternatively, using other software such as QuPath or finetuning a Cellpose model on the specific datasets can increase the accuracy of cell detection.

### Ganglia model

The ganglia segmentation is based on the co-labeling of Hu with another neuronal/glial fiber marker. Within GAT, ganglia are not defined by a minimum number of neurons, so all the cells and ganglion-like structures are included for analysis. This is particularly important in neuropathic conditions such as Hirschsprung disease, where the absence and/or relative reduction of enteric neuronal number is used to support its diagnosis (Burns et al. 2016; Heuckeroth 2018; Niesler et al. 2021; Schäppi et al. 2013). The accuracy of the ganglia segmentation model depends on the quality of the neuronal or glial fiber labeling. High background can affect the accuracy of this approach. The DL approach requires two channels (Hu and a ganglia marker) for defining the ganglia, whereas COUNTEN can segment ganglia with just Hu. As the presence of neurons can be used to define the ganglia, GAT has an additional option to customize ganglia detection using only the Hu labelling. Moreover, for challenging cases, segmentation can be performed in QuPath using pixel classification and the custom ROIs imported back into the GAT workflow.

The reality of DL-based approaches is that GAT may not work accurately on image types it has not encountered before. Generating accurate models requires creating more annotated data, which is made possible within GAT. It is designed to be part of the routine analysis, where the user can opt in to save the images and masks or can be done separately on smaller FOVs. The resulting data can be used for finetuning the GAT models and can be submitted to a public repository. Making annotated data publicly available enables the generation of domain specific image analysis software that can benefit the wider ENS research community (Barth et al. 2023).

### Additional features in GAT

#### Calcium Imaging Analysis

The long-term goal for GAT is to have a collection of common image analysis workflows used within the ENS field. In addition to the analysis of immunofluorescence images, GAT has a calcium imaging analysis workflow aimed at extracting normalized calcium responses from cells in wholemount preparations. The workflow has two parts:

- **Image alignment.** The output of calcium imaging experiments is typically timeseries images which may have movement artefacts either due to drift or tissue movement. GAT provides options to align these images using ‘Linear Stack Alignment with SIFT’ (Lowe 2004) and ‘Template matching’ (Tseng et al. 2011) plugins within Fiji. This can be done on single images or on a folder of images.
- **Extraction of normalized traces.** Once the alignment is performed, the time series image stacks can be imported into Fiji, and the regions of interest (ROIs) manually defined using its drawing tools. The analysis can be performed for multiple cell types in the same image and the extracted traces can be normalized to a user-defined baseline.

